# Tissue-specific dysregulation of mitochondrial respiratory capacity and coupling control in colon-26 tumor-induced cachexia

**DOI:** 10.1101/493056

**Authors:** Andy V. Khamoui, Jessica L. Halle, Gabriel S. Pena, Hector G. Paez, Harry B. Rossiter, Nishant P. Visavadiya, Michael A. Whitehurst

**Affiliations:** Department of Exercise Science and Health Promotion, Florida Atlantic University, Boca Raton, FL 33431, USA; Division of Respiratory and Critical Care Physiology and Medicine, Department of Medicine, Los Angeles Biomedical Research Institute at Harbor-UCLA Medical Center, Torrance, CA 90502, USA; Institute for Healthy Aging and Lifespan Studies, Florida Atlantic University, Boca Raton, FL 33431, USA; Cancer Research Group, Florida Atlantic University, Boca Raton, FL 33431, USA

**Keywords:** skeletal muscle atrophy, liver, adipose, cancer cachexia, OXPHOS, high-resolution respirometry

## Abstract

In addition to skeletal muscle dysfunction, recent frameworks describe cancer cachexia as a systemic disease involving remodeling of non-muscle organs such as adipose and liver. Impairment of mitochondrial function is associated with multiple diseases. The tissue-specific control of mitochondrial function in cancer cachexia is not well-defined. This study determined mitochondrial respiratory capacity and coupling control of skeletal muscle, white adipose tissue (WAT), and liver in colon-26 (C26) tumor-induced cachexia. Tissues were collected from PBS-injected weight-stable mice, C26 mice that were weight-stable, and C26 mice with moderate (10% weight loss) and severe cachexia (20% weight loss). WAT showed high non-phosphorylating LEAK respiration and reduced respiratory control ratio (RCR, index of OXPHOS coupling efficiency) during the induction of cachexia. Liver RCR decreased early due to cancer, and further declined with severe cachexia, where Ant2 but not Ucp2 expression was increased. Ant2 also related inversely with RCR in the liver (r=-0.547, p<0.01), suggesting a role for Ant2 in uncoupling of liver OXPHOS. Increased liver cardiolipin occurred during moderate cachexia and remained elevated in severe cachexia, suggesting this early event may also contribute to uncoupling. Impaired skeletal muscle mitochondrial respiration occurred predominantly in severe cachexia. These findings suggest that mitochondrial function is subject to tissue-specific control during cancer cachexia, whereby remodeling in WAT and liver arise early and could contribute to altered energy balance, followed by impaired skeletal muscle respiration. We highlight an underrecognized role of liver mitochondria in cancer cachexia, and suggest mitochondrial function of multiple tissues to be targets for therapeutic intervention.

## Introduction

Approximately half of all cancer patients undergo cachexia, a life-threatening comorbidity of cancer in which tumor-induced metabolic abnormalities contribute to hallmark clinical features such as involuntary weight loss and skeletal muscle atrophy [1, 2]. Despite impairing responsiveness to anti-cancer treatment and accounting for an estimated 20% of cancer-related deaths [3], cachexia continues to be an underrecognized issue in cancer care, and a major source of frustration for patients and family members alike [1, 4]. Because the root causes of cancer cachexia are not well-defined at present, effective treatment options remain elusive [5]. While research efforts often emphasize skeletal muscle pathophysiology, current frameworks describe a systemic condition in which multiple organs such as adipose, bone, brain, heart, and liver are reprogrammed or remodeled to generate the cachectic phenotype [2, 6, 7]. Convincing evidence supports the existence of cross-talk mechanisms between several of these organs, and targeted treatment of non-muscle organs have been shown to rescue losses of body weight and muscle mass [8, 9]. The multi-organ involvement underscores the highly complex nature of cancer cachexia, whereby multiple mechanisms of metabolic disturbance may be responsible for the hallmark clinical manifestations.

Several lines of evidence implicate mitochondria in the pathogenesis of cancer cachexia [10, 11]. Mitochondria are well known for their central role in cellular function due to their regulation of nutrient oxidation and bioenergetics, diverse signaling pathways, and cell fate decisions [12]. Given these critical roles, disturbances to mitochondria and their metabolic functions have been implicated in aging, neurodegenerative disease, and cancer [13-16]. In particular, effects on mitochondrial respiration are important because oxidative phosphorylation (OXPHOS), which couples the electron transfer system (ETS) to ADP phosphorylation, can affect redox status, mitochondrial dynamics, quality control, and hence the overall health of the mitochondrial pool [12]. In cancer cachexia, mitochondrial function has been most widely studied in skeletal muscle, with several mechanisms proposed to link mitochondrial functions to muscle mass. Elevated oxidant emission could lead to protein degradation and muscle atrophy [17, 18]. Further, restricted ATP provision from impaired OXPHOS may cause energetic stress, downstream activation of protein degradation, and muscle dysfunction [18, 19]. Indeed, recent reports found decreased complex I-linked OXPHOS capacity and coupling efficiency *in situ,* and reduced coupling efficiency *in vivo* in skeletal muscle of rodents with cancer cachexia [20-22]. These defects were observed at or near timepoints in which marked cachexia already occurred, and are suggestive of muscle dysfunction secondary to global changes in systemic metabolism [23]. How skeletal muscle respiration might be affected in other defined coupling and substrate states throughout the development of cancer cachexia, from early to late stage, requires further investigation.

In addition to skeletal muscle oxidative metabolism, considerable interest has been devoted to adipose tissue function as a cause of cachexia. White adipose tissue depots have been shown to undergo a phenotypic switch to resemble the more metabolically active, mitochondrial-dense, heat producing brown adipose compartment (i.e. browning) [8, 24]. In other conditions characterized by severe metabolic stress and browning (i.e. burn injury), WAT shows high LEAK respiration [25], which reflects inner membrane leakiness and intrinsic uncoupling, thereby dissipating the proton gradient independent of ATP synthase and generating heat. The metabolic rewiring of WAT has been proposed as a source of elevated energy expenditure and thus involuntary weight loss [24, 25]. It has also been suggested that inefficiency of mitochondrial OXPHOS and uncoupling in the liver could be another mechanism by which energy is dissipated as heat, metabolic rate increases, and weight loss ensues [7, 26]. Although the overall role of liver metabolism in cancer cachexia has been less frequently explored in comparison to skeletal muscle and adipose, given the highly energetic nature of the liver and its control over systemic metabolism, uncoupling of liver OXPHOS represents an attractive hypothesis that should be examined further.

This investigation tested the hypothesis that mitochondrial respiration is subject to tissue-specific control mechanisms during the induction and progression of cancer cachexia, and that these indices of mitochondrial function relate to body weight loss and skeletal muscle atrophy, the hallmark features of cancer cachexia. How mitochondrial respiration functions in skeletal muscle, WAT, and liver before overt features of cachexia occur, along with the extent to which they change as severe cachexia arises, is not known. Using high-resolution respirometry, we conducted a comprehensive analysis of mitochondrial respiratory function during the induction and progression of cancer cachexia in tissues important for energy metabolism and muscle protein turnover, in order to better understand how tissue-specific alterations in mitochondrial performance may contribute to cancer cachexia, whether specific coupling and substrate states are differentially impacted, and to further validate assertions that OXPHOS and mitochondrial function should be considered targets for therapeutic intervention.

## Methods

### Animals and design

Ten week old Balb/c males (Envigo) were randomly assigned to receive either an injection of PBS or colon-26 (C26) tumor cells. The C26 tumor-bearing mouse is a well-established pre-clinical model of cancer cachexia [27-32]. In this model, a tumor growth period of three weeks are typically allowed to elapse before tissue collection in order for salient features of cachexia to develop (i.e. weight loss, muscle atrophy). To evaluate mitochondrial function during the induction and progression of cachexia, tissue was collected from C26 mice at different timepoints after cell injection, followed by stratification into groups according to weight loss in accordance with previous literature [33, 34]. The 4 groups studied included: 1) Tumor-free, weight-stable mice that were PBS injected (PBS-WS, n=4), 2) C26 mice with confirmed tumors that did not exhibit weight loss and were therefore weight-stable (C26-WS, n=6), 3) C26 mice with moderate cachexia (10% weight loss; C26-MOD, n=7), and 4) C26 mice with severe cachexia (≥20% weight loss; C26-SEV, n=6).

These classifications were adapted from prior pre-clinical investigations in which 10% weight loss was considered moderate cachexia, and 20% severe [33, 34]. Weight loss for each mouse was calculated as the percentage change between carcass weight (i.e. tumor-free body weight) and body weight recorded on the day of cell injection. Mice were individually housed, provided food and water *ad libitum,* and maintained on a 12:12 hr light:dark cycle. All procedures were approved by the Institutional Animal Care and Use Committee at Florida Atlantic University (Protocol # A16-39).

### C26 tumor cell culture and injection

C26 cells (CLS Cell Lines Service, Eppelheim, Germany) were cultured in a humidified 5% CO_2_ incubator using complete media that contained RPMI 1640 supplemented with 1% penicillin/streptomycin (vol/vol) and 10% FBS (vol/vol). Media was replaced every two to three days. At sub-confluency, cells were harvested by incubation with trypsin (0.05%, Gibco) and subsequently pelleted by centrifugation. The supernatant was then discarded and the pellet resuspended in sterile PBS. Viable cells were counted in a hemocytometer by trypan blue staining and light microscopy. Mice in C26 groups were gently restrained and injected s.c. in the upper back with a cell suspension containing 1 x 10^6^ cells. Mice assigned to weight-stable control were injected with an equivalent volume of sterile PBS [28, 35].

### Tissue collection and processing

Mice were euthanized by ketamine/xylazine overdose delivered i.p. (300/30 mg/kg). Hindlimb skeletal muscles, vital organs, and epididymal white adipose tissue (WAT) were carefully isolated and removed. The left medial gastrocnemius, left epididymal WAT, and left lateral lobe of the liver, were immediately placed into ice-cold preservation buffer (BIOPS: 2.77 mM CaK_2_EGTA, 7.23 mM K_2_EGTA, 5.77 mM Na_2_ATP, 6.56 mM MgCl_2_·6H_2_O, 20 mM Taurine, 15 mM Na_2_PCr, 20 mM Imidazole, 0.5 mM DTT, 50 mM MES hydrate) and stored on ice in preparation for *in situ* analysis of mitochondrial respiration. The gastrocnemius muscle was selected because it is a major locomotor muscle that undergoes atrophy in this model [32], and has been previously used to investigate mitochondrial respiratory function in mouse studies of metabolic dysfunction [36, 37]. Epididymal WAT was selected due to its anticipated remodeling and relative abundance, which ensured adequate tissue availability for the respirometric assay. Because WAT undergoes depletion to undetectable levels by ~20% weight loss, WAT was not analyzed in the severe cachexia group. The left lateral lobe of the liver was chosen in accordance with Heim et al [38]. The right hind limb muscles were mounted cross-sectionally in tragacanth gum on cork, and frozen in isopentane cooled by liquid nitrogen for later histological analysis. Remaining tissue samples were snap frozen and stored at -80°C.

To prepare WAT for respirometry, a portion of tissue was gently blotted dry, and ~20-50 mg weighed out for each chamber of the respirometer. These WAT samples were then sectioned into 2-3 pieces prior to placement into the chambers. Chemical permeabilization of WAT by addition of digitonin into the chambers was not performed based on preliminary tests and reports by others in which no effect on respiratory capacity was observed [39]. Preparation of the liver for respirometry was adapted from previous work [40]. Briefly, a pair of small sections from the left lateral lobe (~6 mg each) were placed in a petri dish with ice-cold BIOPS and subjected to gentle mechanical separation with forceps under a dissecting microscope. Duplicate liver samples were blotted dry on filter paper, weighed, and placed into the respirometer chambers. To prepare skeletal muscle for respirometry, the gastrocnemius was placed in a petri dish containing ice-cold BIOPS and mechanically separated with sharp forceps into duplicate fiber bundles (~4-6 mg each) under a dissecting microscope [41, 42]. Fiber bundles were then permeabilized by placing them into separate wells of a 6-well plate filled with BIOPS containing saponin (50 μg/ml) and incubated with gentle shaking on ice for 20 minutes. Following saponin treatment, fiber bundles were washed in respiration medium (MiR05) on ice with gentle shaking for 10 min (MiR05: 0.5 mM EGTA, 3 mM MgCl_2_, 60 mM K-lactobionate, 20 mM taurine, 10 mM KH_2_PO_4_, 20 mM HEPES, 110 mM Sucrose, and 1g/l BSA, pH 7.1). After washing, the fiber bundles were blotted dry on filter paper and weighed before being placed into the respirometer chambers.

### High-resolution respirometry

*In situ* respiration was assayed in the pre-determined order of WAT, liver, and skeletal muscle. This sequence was based on reported stability of mitochondrial performance following storage in BIOPS, with skeletal muscle showing the greatest retention of respiratory function whereas WAT shows a more rapid decline [39]. Oxygen flux per tissue mass (pmol·s^-1^·mg-^1^) was recorded in real-time at 37°C in the oxygen concentration range of 550-350 nmol/ml using high-resolution respirometry (Oxygraph-2k, Oroboros Instruments, Innsbruck, AT) and Datlab software (Oroboros Instruments, Innsbruck, AT). In WAT and liver, respiration was assessed by a substrate-uncoupler-inhibitor-titration (SUIT) protocol adapted from Porter et al. [43, 44] containing the following sequential injections: 1) 1 mM malate, 75 μM palmitoyl-carnitine, 5 mM pyruvate, and 10 mM glutamate, to determine non-phosphorylating LEAK respiration supported by complex I linked substrates (with fatty acids) (CI_*L*_); 2) 5 mM ADP to achieve maximal phosphorylating respiration from electron input through complex I (CI_*P*_); 3) 10 mM succinate to saturate complex II and achieve maximal convergent electron flux through complex I and II (CI+II_*P*_); 4) 10 μM cytochrome c to assess the integrity of the outer mitochondrial membrane and hence quality of sample preparation (duplicate samples were rejected when flux increased by >15% [45]); 5) 0.5 μM carbonylcyanide m-cholorophenyl hydrazone (CCCP) to assess complex I and II linked ETS capacity (i.e. maximal capacity of the electron transfer system; CI+II_*E*_); 6) 0.5 μM rotenone to inhibit complex I (CII_*E*_); and 7) 2.5 μM Antimycin A to inhibit complex III and obtain residual oxygen consumption. We note that in our evaluation of OXPHOS supported by complex I and II linked substrates (i.e. CI_*P*_, CI+II_*P*_), a fatty acid was included in the protocol, therefore, electrons are also supplied into the respiratory chain via electron-transferring flavoprotein [42].

For mitochondrial respiration in skeletal muscle, a similar SUIT protocol was followed with slight modifications to the sequence of injections in order to determine fatty acid based respiration [46]: 1) 1 mM malate and 75 μM palmitoyl-carnitine to determine LEAK respiration supported by fatty acids (FAO_*L*_); 2) 5 mM ADP to determine fatty acid OXPHOS capacity (FAO_*P*_); and 3) 5 mM pyruvate and 10mM glutamate to evaluate complex I supported OXPHOS capacity (with fatty acids) (CI_*P*_). Subsequent assessment of CI+II_*P*_, outer membrane integrity, CI+II_*E*_, CII_*E*_, and residual oxygen consumption were identical to steps 3-7 of the protocol used for WAT and liver.

### Data reduction and analysis

Oxygen fluxes of the different respiratory states were corrected by subtracting the residual oxygen consumption. Fluxes from each duplicate measurement were averaged for statistical analysis. To determine flux control ratios, which express respiratory control independent of mitochondrial content, tissue mass-specific oxygen fluxes from the SUIT protocol were divided by maximal electron transfer system capacity (CI+II_*E*_) as the reference state [42]. Because CI+II_*E*_ is an intrinsic indicator of mitochondrial function that represents the maximal capacity of the electron transfer system, it can be used to normalize the other respiratory states [42]. The respiratory control ratio (RCR), an index of coupling efficiency of the OXPHOS system, was calculated for WAT and liver in the complex I linked substrate state by dividing CI_*P*_ and CI_*L*_ *(P/L)* [47]. The inverse RCR in the complex I supported state *(L/P)* was also calculated. To determine the fraction of maximal OXPHOS capacity serving LEAK respiration, the oxygen flux measured with complex I substrates but not adenylates, CI_*L*_, was divided by CI+II_*P*_. [42, 47] The substrate control ratio (SCR), which evaluates the change in oxygen flux by addition of substrate within a defined coupling state, was calculated for succinate (SCR_succinate_) as CI+II_*P*_/CI_*P*_ [44].

### Total homogenate and subcellular fractionation

Tissue homogenate (skeletal muscle, WAT, and liver) was prepared using a Potter-Elvehjem homogenizer containing 1 mL of ice-cold mitochondrial isolation buffer (215 mM mannitol, 75 mM sucrose, 0.1% BSA, 20 mM HEPES, 1 mM EGTA, and pH adjusted to 7.2 with KOH), as previously described [48]. 400 μl of tissue homogenate was frozen immediately in −80 °C for biochemical assays. To isolate the mitochondria, the remaining tissue homogenate was centrifuged at 1,300 g for 3 min at 4 °C to obtain nuclear pellets. The supernatant was further centrifuged at 10,000 g for 10 min at 4 °C to obtain mitochondrial pellets. The final mitochondrial pellet was resuspended in 40 μl isolation buffer. The protein content of total homogenate and mitochondrial fraction was measured using the BCA protein assay kit.

### H_2_O_2_ production

Liver and skeletal muscle mitochondrial H_2_O_2_ production were measured using 50 μM Amplex Red (Cat#10187-606, BioVision) and 1 U/ml *horseradish peroxidase* (HRP) reagents at 30 °C as described previously [49]. The formation of fluorescent resorufin from Amplex red was measured after a 10 min period at 530-nm excitation and 590-nm emission filters using a Biotek Synergy HTX spectrofluorometer (Winooski, VT).

### Citrate synthase activity

Citrate synthase (CS) was analyzed as a surrogate for mitochondrial content in homogenized liver and gastrocnemius tissues. CS activity was determined using a commercially available kit according to the manufacturer’s instructions (MitoCheck^®^ Citrate Synthase Activity Assay Kit, Cayman Chemical). Absorbance was measured spectrophotometrically in 30 second intervals for 20 minutes at 412 nm. Samples were analyzed in duplicate at a tissue concentration of 2 mg/ml. CS activity was expressed as nM/min/μg protein.

### Cardiolipin content

The fluorescent dye 10-N-Nonyl-Acridine Orange (Cat # A7847, Sigma) was used specifically to measure mitochondrial cardiolipin content [50, 51]. Briefly, 50 μg of liver mitochondria and white adipocyte total protein homogenate was incubated with 50 μM of NAO reagent in mitochondrial isolation buffer at 30°C for 20 min in the dark. After incubation, the red fluorescence of NAO bound to cardiolipin was measured at wavelengths of 495 nm (excitation) and 519 nm (emission) with a Biotek spectrofluorometer.

### Myofiber cross-sectional area

Procedures for determining myofiber cross-sectional area (CSA) were performed as previously described [32]. Briefly, transverse sections 8 μm thick were sectioned from the mid-belly of the gastrocnemius on a cryostat at -20°C. Sections were subsequently fixed with 10% formalin, stained with hematoxylin, washed with PBS, and coverslipped with Immu-Mount medium. Images were acquired at 20x and analyzed by NIH-Image J software.

### Western blotting

A total of 30 μg of protein from total homogenate or mitochondrial fraction of liver and skeletal muscle were resolved by SDS-PAGE using 4–20% Criterion™ TGX™ Precast gels (Cat# 5671095, Bio-Rad, Hercules, CA). The proteins were transferred onto polyvinylidene difluoride (PVDF) membranes, blocked with 6% nonfat dry milk or 5% BSA (for phospho-specific antibodies) for one hour at room temperature, and then incubated at 4°C overnight with the primary antibody of interest. The primary antibodies included: mitochondrial anti-adenine nucleotide translocase 2 rabbit mAb (Ant2, 1:2500 dilution, cat#14671), mitochondrial anti-uncoupling protein 2 rabbit mAb (Ucp2, 1:2500 dilution, cat#89326), anti-phospho-AMP activated protein kinase α rabbit mAb (Thr172) (p-AMPKα, 1:2000, cat#2535), total anti-AMP activated protein kinase α rabbit polyAb (AMPKa, 1:2000 dilution, cat#2532), anti-α-tubulin mouse mAb (1:5000, cat#3873) and mitochondrial anti-voltage dependent anion channel rabbit mAb (VDAC, 1:5000 dilution, cat#4661) from Cell Signaling Technology Inc. The mitochondrial anti-creatine kinase 2 rabbit polyAb (CKMT2, 1:3000 dilution, cat#SAB2100437) was from Sigma-Aldrich. For secondary antibodies, peroxidase-conjugated horse anti-mouse IgG (cat#7076) and goat anti-rabbit IgG (cat # 7074) were obtained from Cell Signaling Technology. The immunoreactive protein reaction was revealed using SuperSignal™ West Pico PLUS Chemiluminescent Substrate (cat# PI34580, Thermo Fisher). The reactive bands were detected by ChemiDoc™ XRS+ imaging system (Bio-rad) and density measured using NIH ImageJ software.

### Statistical analysis

All data are reported as mean±SE. Group differences were determined by one-way ANOVA. In the event of a significant F-test, posthoc analysis was conducted using Tukey’s HSD. Pearson correlation coefficients (r) were used to determine the relationship between indices of mitochondrial respiration, percent body weight change, and myofiber cross-sectional area. Pearson r correlation coefficients were also used to determine the relationship between mitochondrial respiration and expression of selected proteins. Level of significance was accepted at p<0.05.

## Results

### Weight loss and organ atrophy in colon-26 tumor-induced cachexia

Body weight change averaged -10% in C26-MOD, and -22% in C26-SEV, which were significantly different from PBS-WS and C26-WS (Fig. 1a). Weight loss was also significantly greater in C26-SEV compared to C26-MOD (Fig. 1a). Anorexia was not present as mean food intake per day was not different between the C26 groups and PBS-WS (p>0.05) (data not shown). Tumor burden increased in accordance with weight loss (Fig. 1b). Muscle weights were ~20-30% lower in C26-MOD and C26-SEV compared to PBS-WS and C26-WS (Fig. 1c). Epididymal fat was substantially depleted in C26-MOD relative to PBS-WS and C26-WS; epididymal fat was not detected in C26-SEV. (Fig. 1d). In comparison to PBS-WS, the spleen was enlarged in all C26 groups (+34-74%) (Fig. 1e), indicating an inflammatory response to tumor load. Absolute liver mass was lower in C26-MOD and C26-SEV compared to the WS groups (Fig. 1f), but when adjusted for body weight no group differences were observed (Fig. 1g). Fiber cross-sectional area was ~45% lower in C26-MOD, and ~55% lower in C26-SEV compared to the WS groups (Fig. 1h-i). Fiber size distribution revealed the greatest percentage of small fibers in C26-SEV, followed by C26-MOD (Fig. 1j).

**Figure 1.**
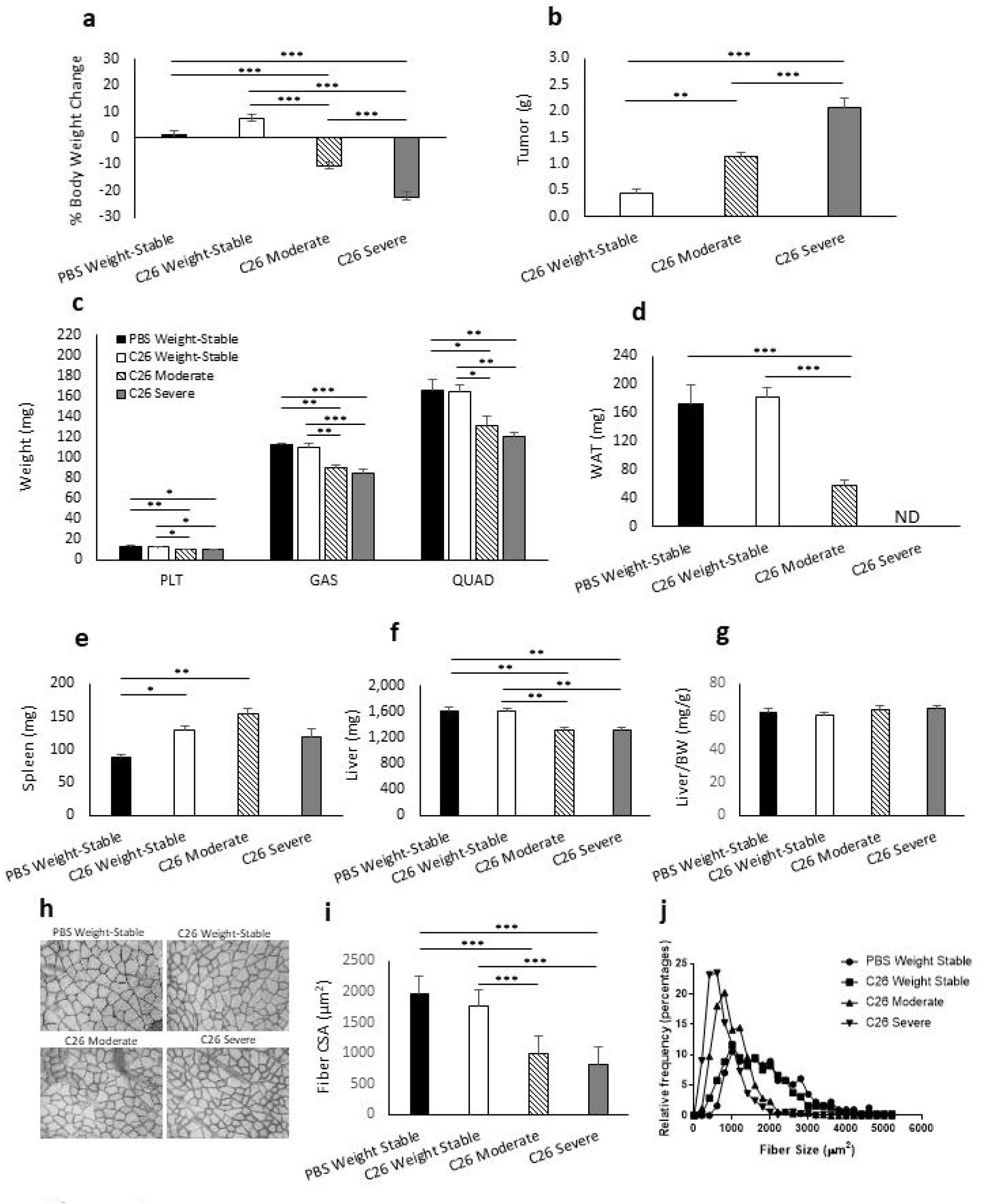
Weight loss and organ atrophy in colon-26 tumor-induced cachexia. (a) Body weight changes of PBS Weight-Stable (n=4), C26 Weight-Stable (n=6), C26 Moderate (n=7), and C26 Severe (n=6). (b) Tumor weights of experimental groups. (c) Skeletal muscle wet weights of plantaris (PLT), gastrocnemius (GAS), and quadriceps (QUAD). (d) Epididymal white adipose tissue (WAT) wet weight. WAT was not detected (ND) in C26 severe group. (e-g) Wet weight of spleen (e) and liver (f), and liver weight normalized to body weight (g). (h) Representative myofiber cross-sections of gastrocnemius muscle imaged at 20x. Imaged cross-sections were analyzed for all mice excluding n=1 from C26 Weight-Stable, and n=2 from C26 Severe due to unavailable tissue mounts (total analyzed n=20). (i) Mean myofiber cross-sectional area. (j) Fiber size distribution between groups displayed as relative frequency (percentage). Data presented as mean ± SE. Differences determined by one-way ANOVA. p<0.05 (*), p<0.01 (**), p<0.001 (***).

### Impairment of complex I-supported skeletal muscle mitochondrial respiration in late cachexia

In comparison to PBS-WS and C26-WS, mass-specific fluxes for FAO_*L*_, FAO_*P*_, CI_*P*_, CI+II_*P*_, and CI+II_*E*_ were lower in C26-MOD and C26-SEV (Fig. 2a), indicating a general cachexia-associated loss of muscle respiratory capacity per tissue mass under various substrate and coupling states. In particular, CI_*P*_, CI+II_*P*_, and CI+II_*E*_ were 23-40% lower in C26-MOD, and 58-79% lower in C26-SEV (Fig. 2a). Oxygen fluxes for select substrate and coupling states were also lower in C26-SEV compared to C26-MOD (Fig. 2a), supportive of a progressive deterioration in muscle mitochondrial function per tissue mass as cachexia worsens. There were no differences in CS activity (p>0.05) (Fig. 2d), therefore the impairment of skeletal muscle respiration was not due to lower mitochondrial content. Mass-specific fluxes FAO_*L*_, FAO_*P*_, CI_*P*_, CI+II_*P*_, and CI+II_*E*_ all related linearly with body weight change (r=0.689-0.804) and fiber CSA (r=0.684-0.762) (Figs. 3a-e, h-l).

**Figure 2.**
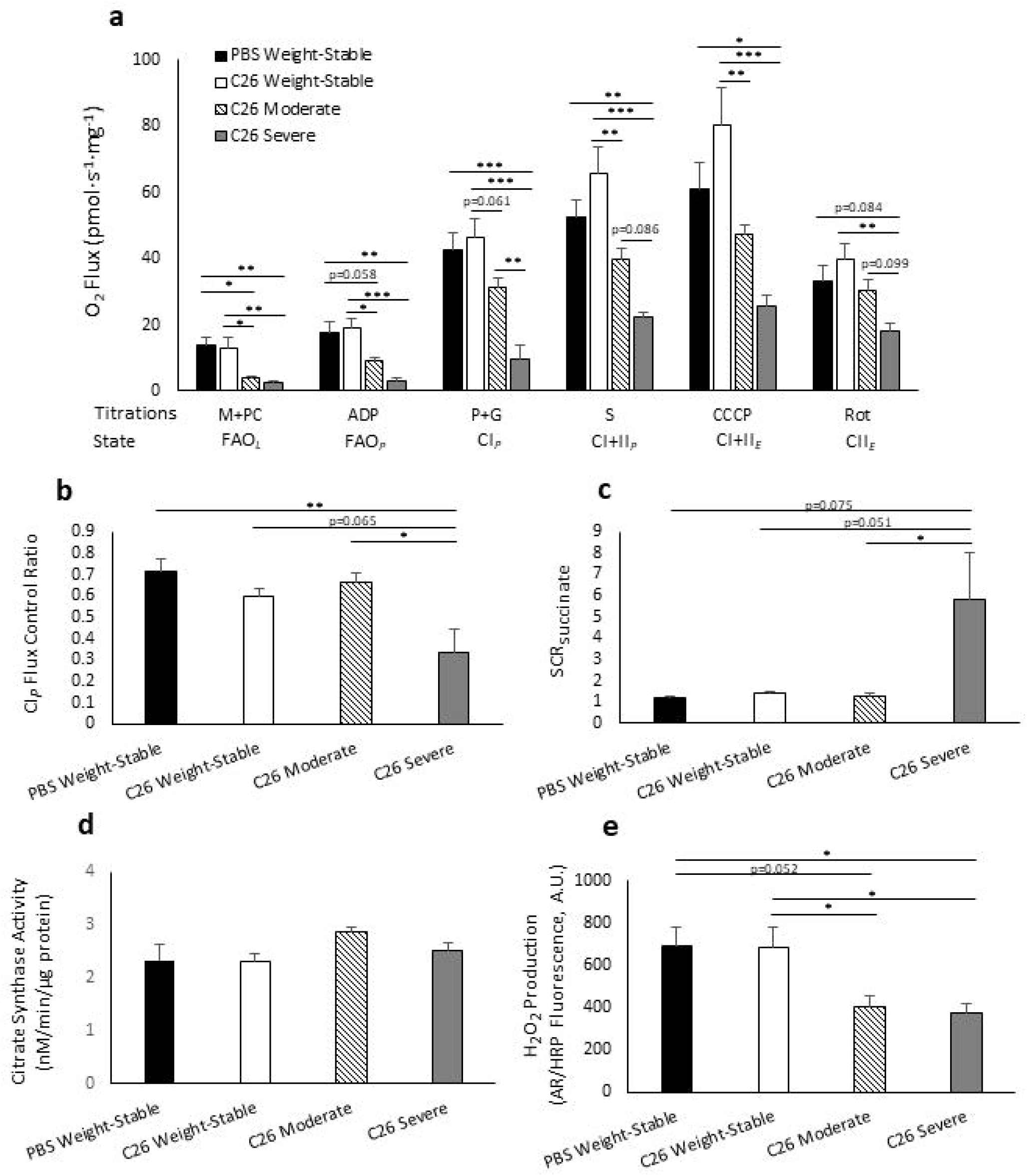
Impairment of complex I-linked skeletal muscle mitochondrial respiration in severe cachexia. (a) Mass-specific oxygen (O_2_) flux of gastrocnemius muscle determined *in situ* by a substrate-uncoupler-inhibitor titration protocol, including fatty acid supported LEAK (FAO_*L*_) through addition of malate and palmitoyl-carnitine (M+PC); fatty acid supported oxidative phosphorylation (OXPHOS) (FAO_*P*_) by addition of adenosine diphosphate (ADP); complex I supported OXPHOS (CI_*P*_) by addition of pyruvate and glutamate (P+G); complex I+II supported OXPHOS (CI+II_*P*_) by addition of succinate (S); maximal electron transfer system (ETS) capacity (CI+II_*E*_) by stepwise addition of carbonyl cyanide m-chlorophenyl hydrazine (CCCP); and complex II ETS (CII_*E*_) by addition of rotenone (Rot). (b) Flux control ratio for complex I supported OXPHOS (CI_*P*_/CI+II_*E*_). (c) Substrate control ratio (SCR) for succinate calculated by dividing CI+II_*P*_ by CI_*P*_. (d) Citrate synthase enzyme activity in gastrocnemius muscle homogenate. (e) Hydrogen peroxide (H_2_O_2_) production in quadriceps muscle mitochondria. Data presented as mean ± SE. Tissues assayed from PBS Weight-Stable (n=4), C26 Weight-Stable (n=6), C26 Moderate (n=7), and C26 Severe (n=6). Differences determined by one-way ANOVA. p<0.05 (*), p<0.01 (**), p<0.001 (***).

Normalization of mass-specific fluxes to ETS capacity, an internal mitochondrial marker, yielded flux control ratios that are independent of mitochondrial density, therefore providing an index of mitochondrial quality. The flux control ratio for FAO_*P*_ (FAO_*P*_/CI+II_*E*_) was 33% lower in C26-MOD (p=0.058), and 60% lower in C26-SEV (p=0.001) compared to PBS-WS (data not shown). FAO_*P*_/CI+II_*E*_ in C26-SEV was also lower than C26-WS (-51%, p=0.007) and C26-MOD (-40%, p=0.086) (data not shown). Suppression of fatty acid based respiration may therefore depend on cachexia severity. The flux control ratio for CI_*P*_ (CI_*P*_/CI+II_*E*_) was 44-53% lower in C26-SEV compared to PBS-WS, C26-WS, and C26-MOD (Fig. 2b). The lower ETS-normalized CI_*P*_ in C26-SEV suggests that muscle mitochondrial quality was impaired, primarily as a consequence of severe, late stage cachexia, and that the source of this dysfunction may reside at complex I. The substrate control ratio for succinate, SCR_succinate_, was ~4-5-fold greater in C26-SEV compared to PBS-WS, C26-WS, and C26-MOD (Fig. 2c). This may implicate a compensatory reliance of severely cachectic muscle on electron supply through complex II in order to stimulate OXPHOS. Body weight change related significantly with both CI_*P*_/CI+II_*E*_ (r=0.487) and SCR_Succinate_ (r=-0.476) (Fig. 3f-g).

**Figure 3.**
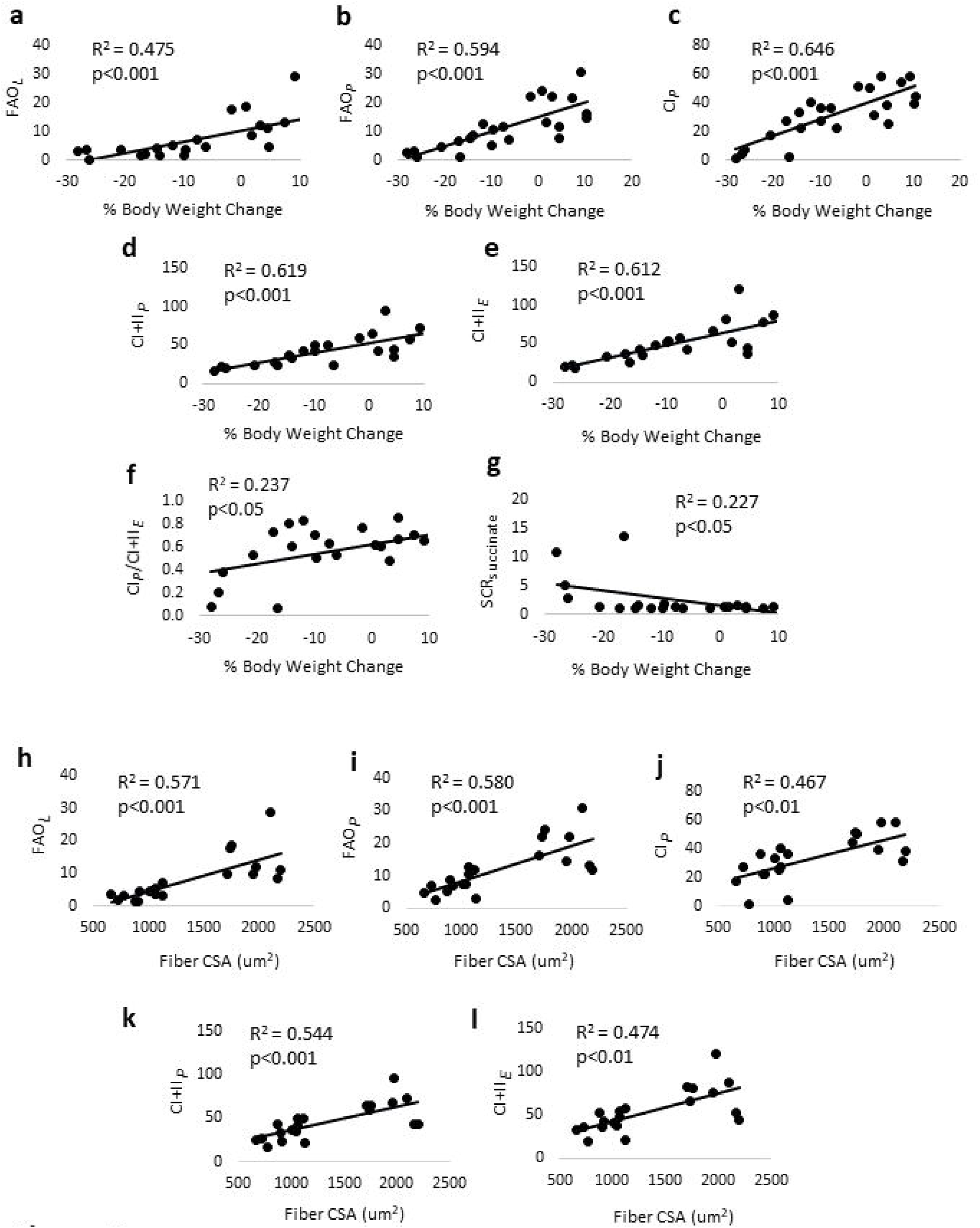
Association of skeletal muscle mitochondrial respiration with weight change and myofiber size. Pearson correlation coefficients were determined for percent body weight change and (a) Fatty acid oxidation LEAK state (FAO_*L*_); (b) Fatty acid oxidative phosphorylation (OXPHOS) (FAO_*P*_); (c) Complex I supported OXPHOS (CI_*P*_); (d) Maximal OXPHOS with electron input through complex I+II (CI+II_*P*_); (e) Maximal electron transfer system (ETS) capacity with electron input through complex I+II (CI+II_*E*_); (f) Complex I supported OXPHOS (CI_*P*_) normalized to maximal ETS capacity (CI+II_*E*_); and (g) Substrate control ratio (SCR) for succinate calculated by dividing CI+II_*P*_ by CI_*P*_. (h-l) Pearson correlation coefficients were also determined for fiber cross-sectional area (CSA) and the same mass-specific oxygen fluxes described in a-e.

H_2_O_2_ in the mitochondrial fraction of skeletal muscle was ~40-50% lower in C26-MOD and C26-SEV compared to PBS-WS and C26-WS (Fig. 2e). Phosphorylation of AMPK was ~100-150% greater in C26-WS and C26-MOD compared to PBS-WS (Fig. 4a, c), indicating early activation of AMPK in skeletal muscle. CKMT2 expression in C26-MOD was ~4-fold greater than PBS-WS, and ~2-fold greater than C26-WS (p=0.091) and C26-SEV (p=0.054) (Fig. 4b, d), consistent with energetically stressed skeletal muscle in early cachexia. Ant2 expression showed a similar pattern of response to CKMT2 (Fig. 4b, e).

**Figure 4.**
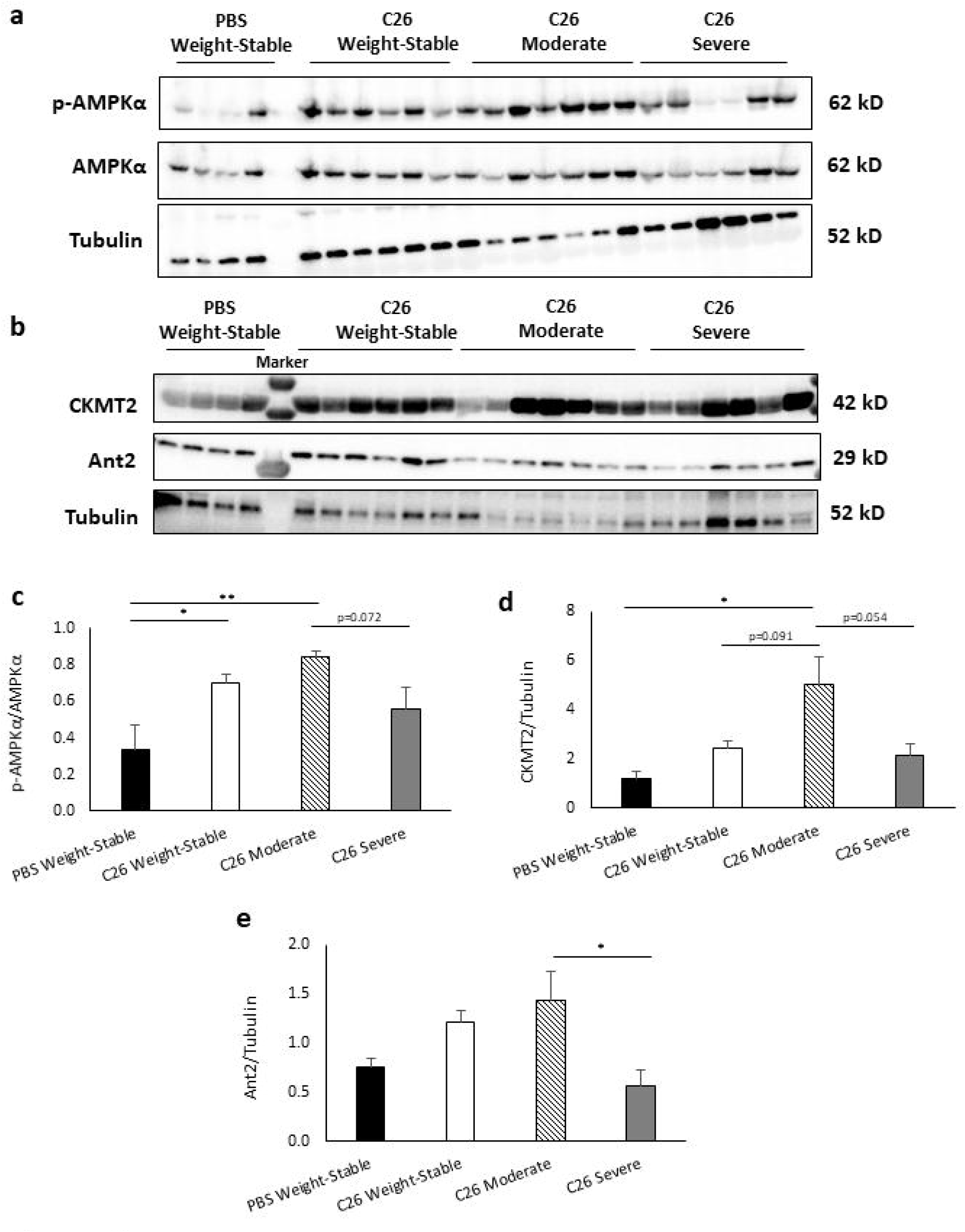
AMPK activation and ADP transport proteins in skeletal muscle of colon-26 mice. (a-b) Immunoblots for p-AMPKa, AMPKa, mitochondrial creatine kinase (CKMT2), Ant2, and tubulin in skeletal muscle homogenate. (c) p-AMPKa expression normalized to total AMPKa. (d) CKMT2 normalized to tubulin. (e) Ant2 normalized to tubulin. Data presented as mean ± SE. Tissues assayed from PBS Weight-Stable (n=4), C26 Weight-Stable (n=6), and C26 Moderate (n=7). Differences determined by one-way ANOVA. p<0.05 (*), p<0.01 (**).

### Increased respiratory rates and uncoupling in WAT during the induction of cancer cachexia

Mass-specific fluxes for WAT including CI_*L*_, CI_*P*_, CI+II_*P*_, and CI+II_*E*_ were significantly greater in C26-MOD compared to the WS groups (Fig. 5a), pointing to an overall increase in metabolic rate. CI_*L*_ in particular showed robust expansion, exceeding the WS groups by ~200% (Fig. 5a). This indicates an increased rate of non-phosphorylating LEAK respiration, and reflects greater leakiness of the inner membrane. RCR for WAT was ~50% lower in C26-MOD compared to the WS groups (Fig. 5b), suggesting loss of OXPHOS coupling efficiency. The LEAK to OXPHOS ratios *L/P* and CI_*L*_/CI+II_*P*_ were greater in C26-MOD by 85-94% and 55-86% respectively, compared to the WS groups (Figs. 5c-d). Flux control ratio for CI_*L*_ (CI_*L*_/CI+II_*E*_) was also greater in C26-MOD, by 47-75%, compared to the WS groups (Fig. 5e). These elevated LEAK ratios are consistent with uncoupled mitochondria. Cardiolipin content was ~50% lower in C26-MOD compared to C26-WS (Fig. 5f). CI_*L*_, CI+II_*P*_, and CI+II_*E*_ were inversely related to body weight change (r=-0.772-0.834) and fiber CSA (r=-0.732-.817) (Figs. 6a-c, g-i), indicating elevated WAT metabolism in weight-losing mice with smaller myofibers, and lower WAT metabolism in weight-stable mice with larger myofibers. Further, RCR for WAT was positively associated with body weight change and fiber CSA (r=0.709, r=0.565) (Fig. 6d, j), whereas the LEAK ratios *L/P* and CI_*L*_/CI+II_*P*_ related inversely with body weight change and fiber CSA (r=-0.628-0.789) (Fig. 6e-f, k-l), suggesting uncoupled WAT mitochondria to be a feature of tumor-induced weight loss and myofiber atrophy.

**Figure 5.**
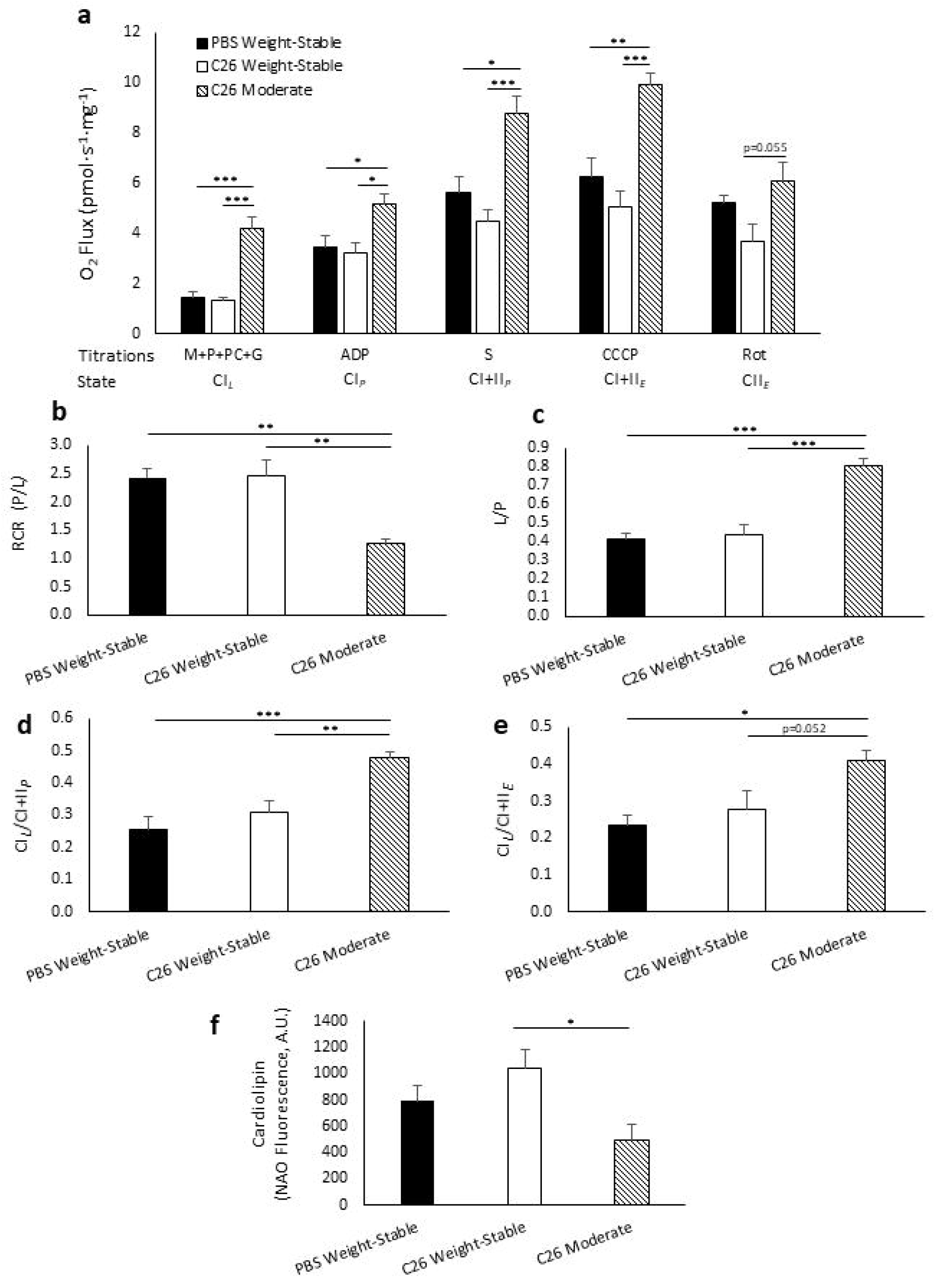
Increased respiratory rates and uncoupling in white adipose tissue (WAT). (a) Mass-specific oxygen (O_2_) flux of WAT determined *in situ* by a substrate-uncoupler-inhibitor titration protocol, including complex I supported LEAK (CI_*L*_) through addition of malate (M), pyruvate (P), palmitoyl-carnitine (PC), and glutamate (G); complex I supported oxidative phosphorylation (OXPHOS) (CI_*P*_) by addition of adenosine diphosphate (ADP); complex I+II supported OXPHOS (CI+II_*P*_) by addition of succinate (S); maximal electron transfer system (ETS) capacity (CI+II_*E*_) by stepwise addition of carbonyl cyanide m-chlorophenyl hydrazine (CCCP); and complex II ETS (CII_*E*_) by addition of rotenone (Rot). (b) Respiratory control ratio (RCR) determined by dividing CI_*P*_ by CI_*L*_. (c) *L/P* determined by dividing CI_*L*_ by CI_*P*_. (d) Ratio of CI_*L*_ and maximal OXPHOS (CI+II_*P*_). (e) Ratio of CI_*L*_ and maximal ETS capacity (CI+II_*E*_). (f) Cardiolipin content in WAT homogenate. Data presented as mean ± SE. Tissues assayed from PBS Weight-Stable (n=4), C26 Weight-Stable (n=6), and C26 Moderate (n=7). Epididymal WAT was not detected in C26 Severe and thus unavailable for analysis. Differences determined by one-way ANOVA. p<0.05 (*), p<0.01 (**), p<0.001 (***).

**Figure 6.**
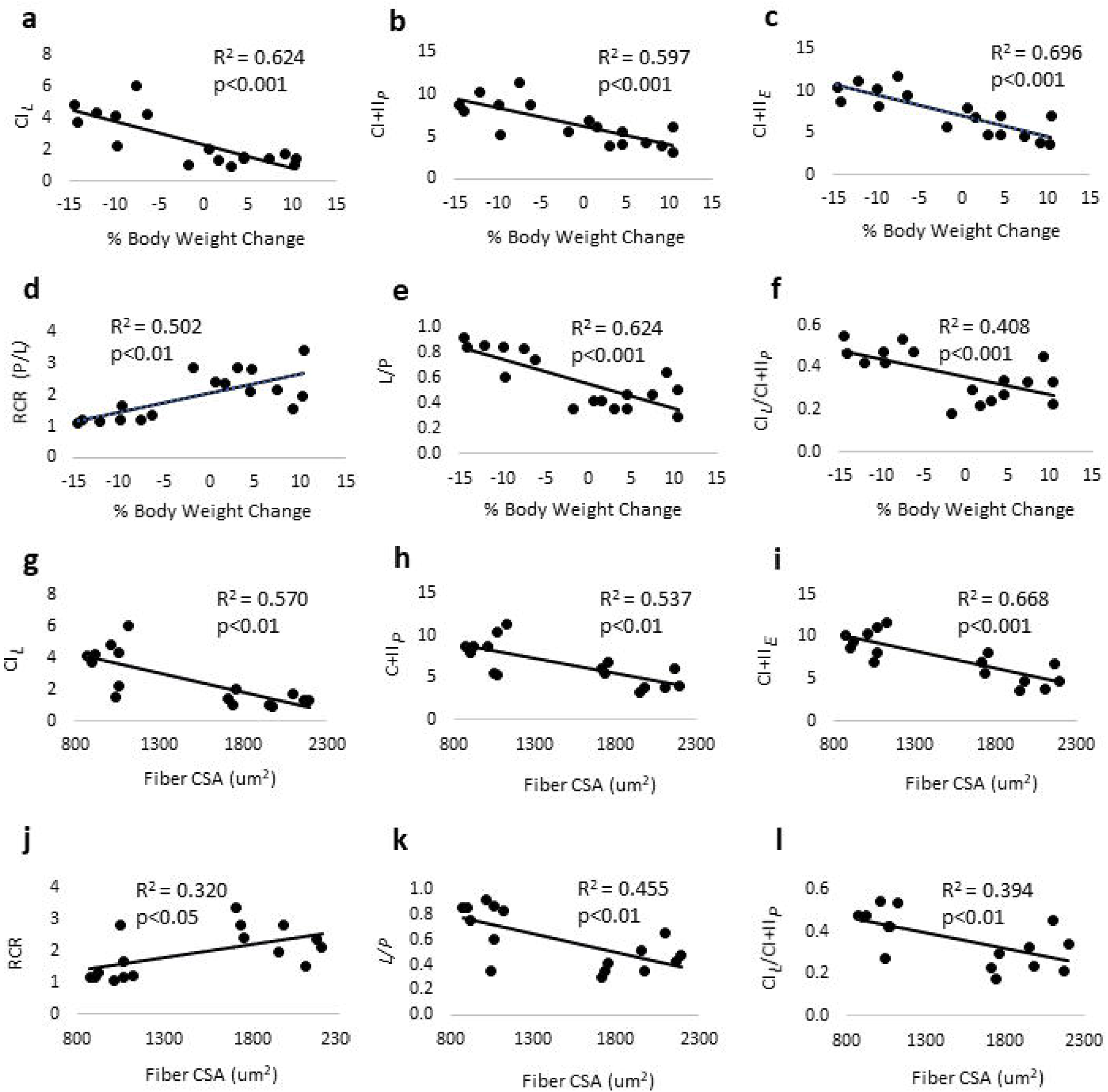
Association of white adipose mitochondrial respiration with weight change and myofiber size. Pearson correlation coefficients were determined for percent body weight change and (a) Complex I supported LEAK (CI_*L*_); (b) Maximal oxidative phosphorylation (OXPHOS) with electron input through complex I+II (CI+II_*P*_); (c) Maximal electron transfer system (ETS) capacity with electron input through complex I+II (CI+II_*E*_); (d) Respiratory control ratio (RCR), calculated by dividing complex I supported OXPHOS (CI_*P*_) by complex I supported LEAK (CIl); (e) *L/P,* calculated by dividing CI_*L*_ by CI_*P*_; and (f) Ratio between CI_*L*_ and CI+II_*P*_. (g-l) Pearson correlation coefficients were also determined for fiber cross-sectional area (CSA) and the same respiratory parameters described in a-f.

### Early loss of liver OXPHOS coupling efficiency and elevated LEAK in C26 mice

Compared to PBS-WS, mass-specific respiration for CI_*L*_, CI_*P*_, CI+II_*P*_, and CI+II_*E*_ were ~40-80% lower in all three C26 groups, indicating early and sustained loss of liver respiratory capacity due to cancer, and not cachexia per se, for each coupling state (i.e. LEAK, OXPHOS, ETS) (Fig. 7a). CS activity, a proxy for mitochondrial density, was not different between groups (p>0.05) (Fig. 7e), suggesting the impairment of liver respiratory function to be independent of mitochondrial mass. In agreement, AMPK phosphorylation status, an upstream signal for PGC-1α-dependent mitochondrial biogenesis, was not different between groups (p>0.05) (Fig. 9b, e). RCR of the liver was ~25-60% lower in C26-WS, C26-MOD, and C26-SEV compared to PBS-WS (Fig. 7b). C26-SEV also had lower liver RCR than C26-MOD (Fig. 7b). Together this may signify an early loss of OXPHOS coupling efficiency due to cancer, that subsequently worsens when severe cachexia develops. CI_*L*_/CI+II_*P*_ was greater in C26-WS (+82%), C26-MOD (+74%), and C26-SEV (+93%) compared to PBS-WS (Fig. 7c), suggesting an early, sustained increase in the fraction of maximal OXPHOS capacity that is LEAK due to cancer rather than cachexia. The *P/E* ratio (CI+II_*P*_/CI+II_*E*_) was greater in C26-SEV compared to all other groups (Fig. 7d). Because *P/E* in C26-SEV approached 1.0 (0.94±0.05), this may indicate dyscoupled liver mitochondria in severe cancer cachexia.

**Figure 7.**
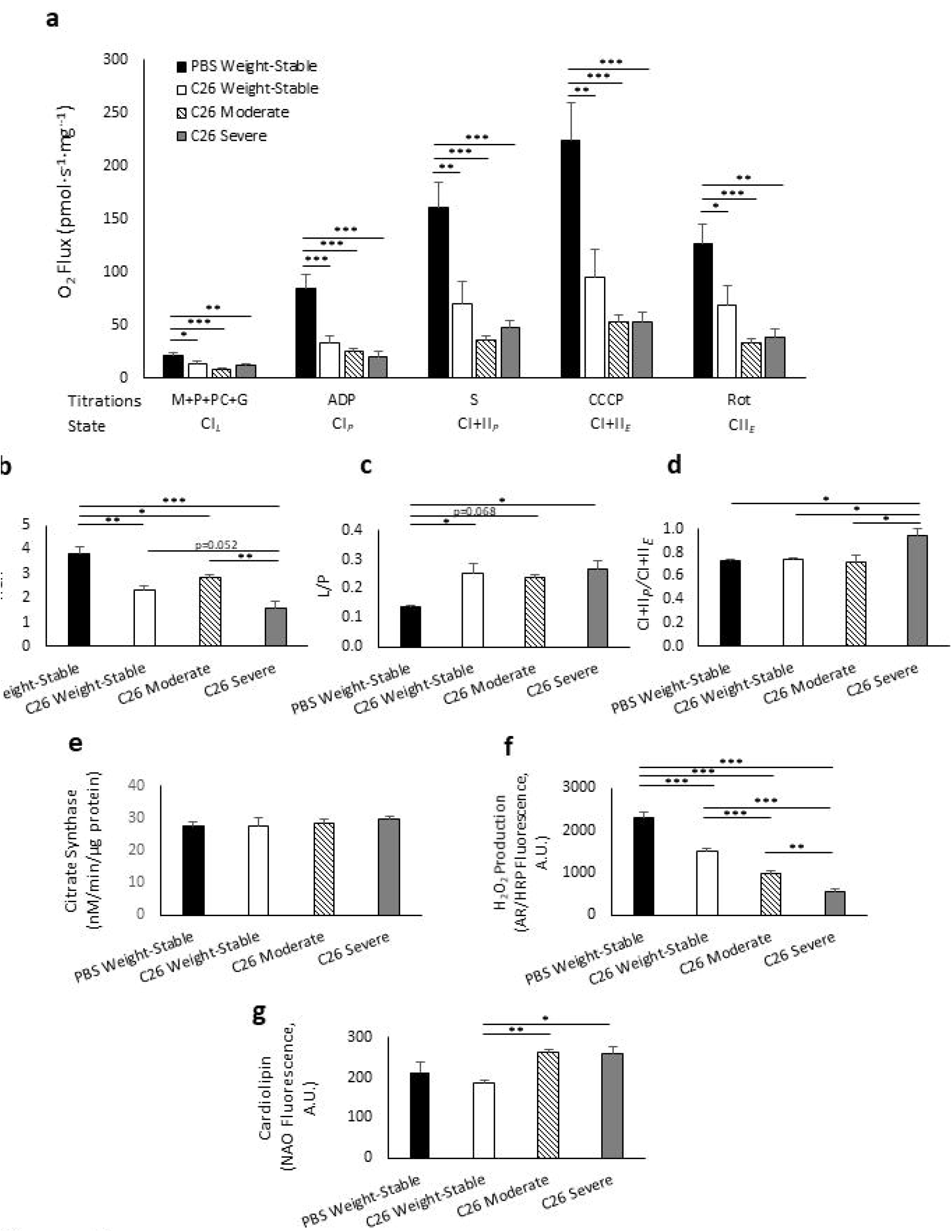
Early loss of liver respiratory function and coupling efficiency in colon-26 mice. (a) Mass-specific oxygen (O_2_) flux of liver measured *in situ* by a substrate-uncoupler-inhibitor titration protocol, including complex I supported LEAK (CI_*L*_) through addition of malate (M), pyruvate (P), palmitoyl-carnitine (PC), and glutamate (G); complex I supported oxidative phosphorylation (OXPHOS) (CI_*P*_) by addition of adenosine diphosphate (ADP); complex I+II supported OXPHOS (CI+II_*P*_) by addition of succinate (S); maximal electron transfer system (ETS) capacity (CI+II_*E*_) by stepwise addition of carbonyl cyanide m-chlorophenyl hydrazine (CCCP); and complex II ETS (CII_*E*_) by addition of rotenone (Rot). (b) Respiratory control ratio (RCR), calculated by dividing CI_*P*_ by CI_*L*_. (c) Ratio between CI_*L*_ and maximal OXPHOS (CI+II_*P*_). (d) The *P/E* ratio, calculated as maximal OXPHOS (CI+II_*P*_) divided by maximal ETS capacity (CI+II_*E*_). (e) Citrate synthase enzyme activity in liver homogenate. (f) Hydrogen peroxide (H_2_O_2_) production in liver mitochondria. (g) Cardiolipin content in liver mitochondria. Data presented as mean ± SE. Tissues assayed from PBS Weight-Stable (n=4), C26 Weight-Stable (n=6), C26 Moderate (n=7), and C26 Severe (n=6). Differences determined by ANOVA. p<0.05 (*), p<0.01 (**), p<0.001 (***).

Mass-specific respiration was positively related to body weight change (r=0.382-0.430) and fiber CSA (r=0.395-0.459) (Figs. 8a-c, g-j), suggesting that cachexia-related weight loss and fiber atrophy may be linked to a depression in liver mitochondrial function. Weight change was also positively associated with liver RCR (r=0.497), but inversely related to the LEAK ratios *L/P* (r=-0.569) and CI_*L*_/CI+II_*E*_ (r=-0.484) (Figs. 8d-f), suggesting that liver mitochondria with tighter coupling appeared more often in weight-stable mice, whereas uncoupling typically appeared with weight loss.

**Figure 8.**
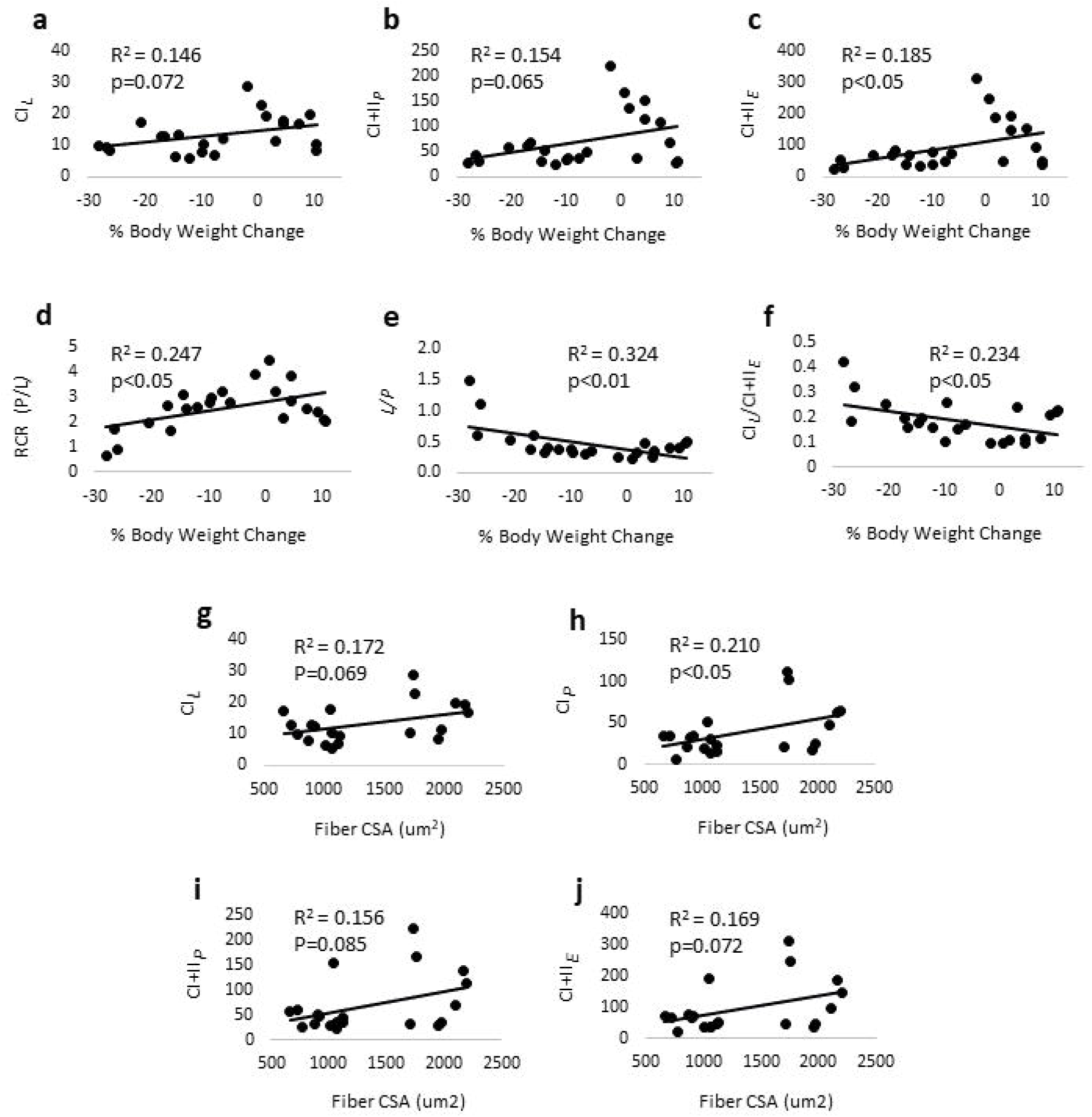
Association of liver mitochondrial respiration with weight change and myofiber size. Pearson correlation coefficients were determined for percent body weight change and (a) Complex I supported LEAK (CI_*L*_); (b) Maximal oxidative phosphorylation (OXPHOS) capacity with electron input through complex I+II (CI+II_*P*_); (c) Maximal electron transfer system (ETS) capacity with electron input through complex I+II (CI+II_*E*_); (d) Respiratory control ratio (RCR), calculated by dividing CI_*P*_ by CI_*L*_; (e) *L/P,* calculated by dividing CI_*L*_ by CI_*P*_. (f) Ratio between CI_*L*_ and CI+II_*E*_. Pearson correlation coefficients were also determined for fiber crosssectional area (CSA) and (g) CI_*L*_; (h) CI_*P*_; (i) CI+II_*P*_; and (j) CI+II_*E*_.

### Decreased ROS, increased cardiolipin, and greater Ant2 expression in cachectic liver mitochondria

To identify events associated with uncoupling of OXPHOS and elevated LEAK, we measured H_2_O_2_ production and cardiolipin content by amplex red and NAO fluorescence, respectively, in the mitochondrial fraction of the liver. H_2_O_2_ is an indicator of mitochondrial ROS emission, and uncoupling may occur as a protective response against high levels of ROS. H_2_O_2_ was significantly lower in all three C26 groups compared to PBS-WS (Fig. 7f). H_2_O_2_ also showed greater decline during weight loss progression, with C26-MOD ~30% lower than C26-WS, and C26-SEV ~40% lower than C26-MOD (Fig. 7f). This gradual decline in ROS emission paralleled decreases in respiratory capacity, and may represent part of a broad, cachexia-associated loss of liver mitochondrial function. Therefore, uncoupling was not due to excessive ROS. Cardiolipin, a phospholipid of the mitochondrial inner membrane that regulates OXPHOS function, was ~40% greater in C26-MOD and C26-SEV compared to C26-WS (Fig. 7g). The greater cardiolipin content in both groups of cachectic mice may support an involvement of this mitochondrial phospholipid in the uncoupling of liver OXPHOS in cancer cachexia. We next probed for Ucp2 and Ant2 expression in liver mitochondria by immunoblotting to determine whether proteins with reported uncoupling properties may be associated with the increased LEAK respiration and uncoupling of OXPHOS (Fig. 9a). Ucp2 protein expression was not significantly different between groups (p>0.05) (Fig. 9a, c). However, Ant2 protein expression was significantly greater in C26-SEV compared to PBS-WS, C26-WS, and C26-MOD, by 30%, 15%, and 16%, respectively (Fig. 9a, d). There was a significant inverse relationship between Ant2 expression and RCR in the liver (r=-0. 547), implying higher liver Ant2 content to be associated with uncoupling of OXPHOS (Fig. 9f).

**Figure 9.**
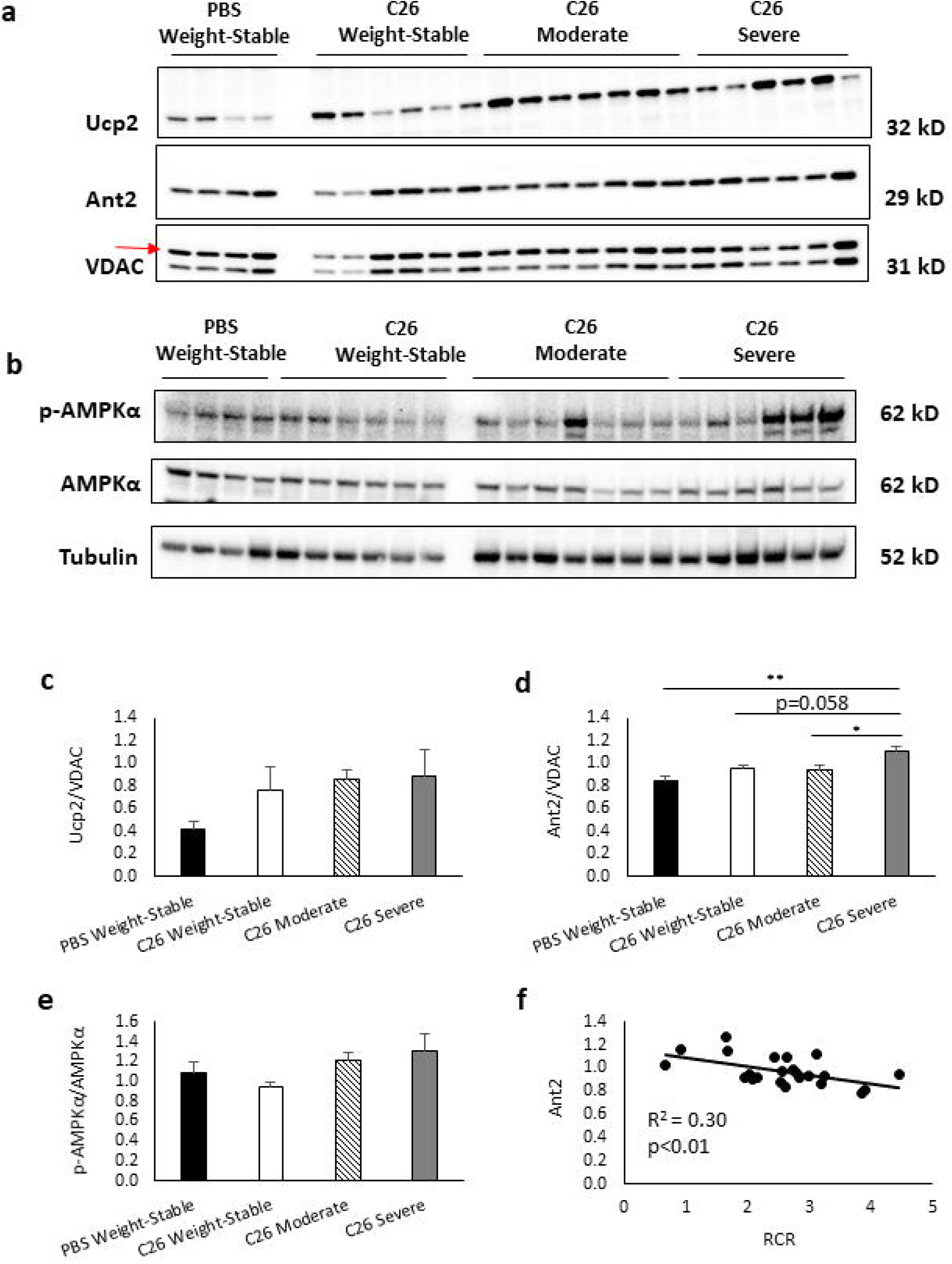
Elevated expression of Ant2 but not Ucp2 in cachectic liver mitochondria of colon-26 mice. (a) Immunoblots for Ucp2, Ant2, and VDAC in mitochondrial lysates from the liver. (b) Immunoblots for p-AMPK, AMPK, and tubulin in liver homogenate. (c) Ucp2 expression normalized to VDAC. (d) Ant2 expression normalized to VDAC. (e) p-AMPK normalized to total AMPK. (f) Association of Ant2 expression with respiratory control ratio (RCR) in the liver. Data presented as mean ± SE. Tissues assayed from PBS Weight-Stable (n=4), C26 Weight-Stable (n=6), C26 Moderate (n=7), and C26 Severe (n=6). Differences determined by one-way ANOVA. p<0.05 (*), p<0.01 (**).

## Discussion

We report tissue-specific alterations in mitochondrial function during colon-26 tumor-induced cachexia. WAT showed high rates of mitochondrial respiration during the induction of cancer cachexia. Non-phosphorylating LEAK respiration of WAT was especially pronounced at this stage. WAT also had reduced efficiency of the OXPHOS system as evidenced by decreased RCR, reflecting uncoupling of respiration from ATP synthesis, possibly due to greater leakiness of the mitochondrial inner membrane. Together these findings are consistent with the presence of uncoupled WAT mitochondria and browning of WAT. RCR of the liver decreased early due to cancer, and further declined with severe cachexia. This happened in concert with increased Ant2 but not Ucp2 expression. Ant2 also exhibited a significant inverse relationship with RCR, suggesting a role for Ant2 in uncoupling of liver OXPHOS in cancer cachexia. Increased liver cardiolipin content occurred during moderate cachexia and remained elevated in severe cachexia, indicating that this may be an early event that contributes to uncoupling of liver OXPHOS. With the exception of fatty acid based respiration, impairment of skeletal muscle mitochondrial respiration was predominant in severe cachexia, and this dysfunction resided at complex I. Mitochondrial respiratory parameters of WAT, liver, and skeletal muscle accounted for a significant proportion of the variance in body weight change and myofiber size (Figs. 3, 6, 8). These findings suggest that mitochondrial function is subject to tissue-specific control during cancer cachexia, whereby early alterations arise in WAT and liver, followed by later impariment of skeletal muscle respiration.

Skeletal muscle has arguably been the most widely studied organ in cancer cachexia. Several pre-clinical investigations have examined *in situ* mitochondrial respiration in skeletal muscle using a cross-sectional design comparing controls and tumor-bearing rodents with marked cachexia. Impaired complex I and complex II OXPHOS were reported [21, 52], consistent with a loss of mitochondrial respiratory function in severely cachectic skeletal muscle. We expand upon these findings by using a SUIT protocol to evaluate *in situ* respiration across a broader range of coupling states and substrate conditions, in tissue obtained from tumor-bearing mice with varying degrees of cachexia. Mass-specific respiration for complex I OXPHOS, and complex I+II OXPHOS and ETS were not significantly different in moderate cachexia compared to PBS weight-stable mice. Significant declines in these variables only occurred in severe cachexia, suggesting that altered skeletal muscle mitochondrial respiration is not an early event in cancer cachexia. Indeed, loss of muscle mass and fiber CSA were evident in mice with moderate cachexia despite no significant impairment of complex I and II linked respiration at this stage. Fatty acid based respiration would be the exception (i.e. FAO_*L*_, FAO_*P*_), which was reduced in moderate cachexia and remained suppressed in the severe state.

Since these parameters represent tissue mass-specific fluxes and do not account for mitochondrial content, we also calculated flux control ratios by normalization to maximal ETS capacity to provide indices of mitochondrial quality that are independent of mitochondrial density. The flux control ratio for complex I OXPHOS in skeletal muscle was impaired only in severe cachexia, consistent with the assertion that muscle mitochondrial dysfunction occurred with late stage cachexia. In line with complex I dysfunction, the substrate control ratio for succinate was ~4-5 fold greater in severe cachexia compared to all other groups, an apparent compensation to stimulate OXPHOS via complex II. Based on the impairment of OXPHOS, we would expect energetic stress to be present in skeletal muscle of cachectic mice. Phosphorylation of AMPK was increased early, in weight-stable C26 mice, and remained elevated in moderate cachexia, which may imply that energetic stress at least partly drives ensuing protein degradation and muscle atrophy. Consistent with energetic stress, mitochondrial creatine kinase (CKMT2) expression was elevated ~2-4-fold in moderate cachexia, which could represent a compensatory mechanism aiming to improve oxidative energy metabolism in skeletal muscle [48]. A similar expression pattern was seen for Ant2, an ADP transport protein, suggesting that Ant2 may also be involved in this compensatory mechanism. In addition to impaired OXPHOS, dysfunctional mitochondria may generate high levels of ROS that in turn affects protein turnover and causes atrophy. We did not observe increased H2O2 in the mitochondrial fraction of skeletal muscle as anticipated. H2O2 production actually declined in moderate and severe cachexia, and does not appear associated with muscle atrophy in this model.

The elevated rates of mitochondrial respiration in WAT were not surprising. A recent report found increased oligomycin and FCCP induced respiration in white adipocytes treated with parathyroid-like protein, a tumor-derived product that induces browning [8]. Others found increased respiratory capacity of brown-like WAT in cachectic mice using glycerol-3-phosphate as a substrate [24]. Our data extends these findings by using a SUIT protocol to evaluate *in situ* respiration of WAT in a broader spectrum of coupling and substrates states. A unique finding was the increase in coupled (phosphorylating) respiration with electron input through complex I and complex I+II, which to our knowledge has not been previously reported. Upregulation of TCA cycle, electron transport, and OXPHOS genes in cancer patients with cachexia has been documented by others [53], and our elevated OXPHOS and ETS capacities are consistent with that gene profile. The increased phosphorylating respiration could be a compensatory response to the lipolysis of WAT. Lipolysis is stimulated to mobilize free fatty acids and glycerol into the bloodstream for use by other tissues. Treatments that inhibit coupling and ATP synthesis are believed to suppress lipogenesis in adipose tissue [54, 55].

Further, insulin-dependent suppression of lipolysis may depend on ATP availability [54, 56]. Thus, increased phosphorylating respiration may be a compensatory effort to provide the energy supply to promote lipogenesis and/or suppress lipolysis, in order to maintain adipose mass in the face of tumor-induced catabolism.

While coupled respiration of WAT increased in early cachexia, the LEAK state showed particularly robust expansion. By dividing coupled and LEAK respiration (in the same complex I-linked substrate conditions), we obtained the respiratory control ratio (RCR), an index of OXPHOS coupling efficiency. Because LEAK expanded to a much larger degree relative to coupled respiration, WAT RCR decreased in early cachexia relative to weight-stable mice. The presence of elevated LEAK was a consistent finding in this study, with greater WAT LEAK in early cachexia even when normalized to maximal OXPHOS and ETS capacities. These findings are significant because they have implications for whole body metabolism. In burn injury, WAT undergoes browning and shows high LEAK respiration in parallel with elevated resting energy expenditure. Reprogramming of WAT metabolism in this manner could potentially contribute to hypermetabolism in cancer cachexia. Therapies which normalize WAT mitochondrial function by reducing LEAK and promoting tighter OXPHOS coupling may be beneficial.

Despite exerting major control over systemic metabolism, the liver has been relatively understudied in cancer cachexia. We observed loss of mass-specific respiration in the liver of all three C26 groups, irrespective of weight loss, suggesting the impairment of respiration to result from tumor load rather than cachexia. RCR was also lower in all three C26 groups, although C26 mice with severe cachexia had the greatest decline. Therefore, the early loss of OXPHOS coupling efficiency arises from tumor load, but subsequently worsens with the development of severe, late-stage cachexia. Consistent with the idea that liver mitochondria are uncoupled in late-stage cachexia, the *P/E* ratio (with an upper limit of 1.0) was greatest in mice with severe cachexia. A *P/E* ratio that approaches 1.0, as was the case in the severely cachectic mice, implies the presence of dyscoupled mitochondria (i.e. pathologically uncoupled) [42] that could be energetically inefficient, and in turn contribute to altered whole body metabolism (e.g. resting energy expenditure).

We are aware of only a few prior investigations on liver mitochondrial energetics in cancer cachexia. Using rats bearing the peritoneal carcinoma as a model of cancer cachexia, Dumas et al. reported reduced OXPHOS efficiency and increased energy wasting in liver mitochondria [26], events in agreement with our finding of reduced liver RCR. They also found the increased energy wasting to coincide with an elevation in cardiolipin content, and coefficient of determination between the two variables indicated substantial shared variance (R^2^=0.64). In the present work, cardiolipin content increased in moderate cachexia, and stayed elevated in the severe state, consistent with their findings. The expansion of cardiolipin mass in liver is of note given that in many pathologies (e.g. heart failure, Barth syndrome, ischemia-reperfusion injury) and aging, modifications to cardiolipin profiles typically consist of decreased content, altered fatty acid composition, and/or peroxidation [57]. We are unable to address whether changes in composition occurred, however the decreased H_2_O_2_ in liver mitochondrial lysates during cachexia suggests minimal peroxidation due to mitochondrial ROS emission. Treatment of normal liver mitochondria with cardiolipin-enriched liposomes has been shown to adversely affect ATP synthesis and increase non-phosphorylating respiration, offering a possible explanation for the significance of enhanced liver cardiolipin in cancer cachexia [58, 59].

The role of Ant in mediating cachexia-associated energy wasting has received some attention [59]. Ant is an ADP/ATP exchanger in the inner mitochondrial membrane that has uncoupling capability. Treatment of liver mitochondria from cachectic rats with carboxyatractylate, an inhibitor of Ant, did not mitigate energy wasting, suggesting that Ant is not the predominant contributor to energetic inefficiency of liver mitochondria from cachectic rodents. The authors therefore concluded that inefficiency of OXPHOS and energy wasting was dependent on cardiolipin, but not Ant. However, we saw increased Ant2 expression in severe cachexia, which would point to an involvement of this molecule in the elevated LEAK state and loss of OXPHOS coupling efficiency. This is supported by the significant inverse relationship between Ant2 and RCR in the liver. Direct manipulations of liver Ant2 are necessary to better understand the role of this molecule in OXPHOS function in cancer cachexia. We also expected to see increased Ucp2 expression in liver mitochondria from C26 mice with cachexia. Although Ucp2 expression was ~2-fold greater in C26 mice compared to PBS-WS, this did not reach statistical significance. Therefore, the increased LEAK respiration and uncoupling of OXPHOS does not appear to be mediated by Ucp2, at least in colon-26 tumor-induced cachexia. These findings bring attention to the previously underappreciated role of liver mitochondrial function in cancer cachexia, and suggest a benefit of therapies that improve mitochondrial function of the liver.

In conclusion, we provide evidence for tissue-specific control of mitochondrial function in colon-26 tumor-induced cachexia. Impairment of skeletal muscle mitochondrial OXPHOS occurred predominantly in severe, late stage cachexia, whereas alterations to WAT and liver arise earlier and could contribute to altered whole body energy balance and involuntary weight loss characteristic of cancer cachexia. Together these findings suggest mitochondrial function of multiple tissues to be potential sites of targeted therapies.

## Acknowledgements

We extend our sincere thanks to Dr. Chun-Jung Huang, Director of the Exercise Biochemistry Laboratory, for research support, and Joseph P. Carzoli and Trevor K. Johnson for technical assistance. JLH was supported by an Undergraduate Research Fellowship (SURF) from the Office of Undergraduate Research and Inquiry at Florida Atlantic University.

